# POI-associated LMD-3 mutation impairs B12-regulated lysosomal function and reproductive capacity in *C. elegans*

**DOI:** 10.1101/2025.10.14.682338

**Authors:** Yile Zhai, Tiantian Wang, Meixi Gong, Yuhan Liang, Wenfei Li, Xin Wang, Zhe Zhang

## Abstract

Reproductive longevity decline is a key feature of aging and premature ovarian insufficiency (POI). While proteostasis collapse is implicated in reproductive aging, the molecular link remains elusive. Here, we engineered the clinically relevant W690C mutation into *C. elegans lmd-3* gene, the human *NCOA7* ortholog, to establish a valuable in vivo POI model. The mutation severely compromises LMD-3 protein stability and expression, leading to a profound collapse in reproductive capacity driven by germline apoptosis. We demonstrate that this defect drives catastrophic autophagic-lysosomal dysfunction, blocking degradation and causing proteotoxic accumulation of misfolded proteins. Crucially, we define this pathology as a functional B12 deficiency and show that mecobalamin (meCbl) supplementation successfully restores proteostasis. Together, these findings delineate a conserved genetic pathway from LMD-3 destabilization to reproductive failure and propose vitamin B12 as a readily translatable therapeutic intervention for age-related reproductive decline.

## Introduction

Reproductive longevity is fundamental for female fertility and healthy aging, yet its decline is one of the first biological processes to emerge in many organisms, from *C. elegans* to humans^1,2^. In humans, the ovary is uniquely susceptible, with its function diminishing decades before other somatic systems^3,4^. This early and pronounced decline contributes significantly to age-related infertility and premature ovarian insufficiency (POI), posing a major global health challenge^1,5^. A growing body of evidence links this decline to impaired proteostasis, the cellular network that maintains protein quality control^6^. While cellular stress responses and protein quality control systems are evolutionarily shared across species^7–9^, the specific molecular pathways that connect these processes to reproductive failure remain elusive.

The nematode *C. elegans* provides a powerful and evolutionarily conserved model for investigating reproductive aging^10,11^. Its distinct and early decline in fertility mirrors the human condition, making it an ideal platform to dissect the molecular basis of reproductive health^2,12^. This decline is largely driven by a loss of germline integrity and function, which is highly dependent on the maintenance of a healthy proteome, a process regulated by conserved cellular systems, including the lysosomal and autophagic pathways^13–15^.

The oxidation resistance 1 (OXR1) protein family is a key player in this process, comprised of OXR1 and nuclear receptor coactivator 7 (NCOA7) in mammals^16^. This evolutionarily conserved family acts as a critical protective mechanism against cellular stress^17,18^. NCOA7 has recently been identified as a key regulator of ovarian longevity in humans and mice, and its orthologs have been shown to protect against oxidative damage in other species like *C. elegans* and zebrafish^19–21^. The TLDc domain of these proteins is known to directly interact with the V-type ATPase, a proton pump essential for lysosomal acidification and degradative function^22,23^. This shared function and evolutionary conservation highlight the potential of using *C. elegans* model to gain insights into human disease.

In this study, we engineered a clinically relevant human mutation in *C. elegans* to investigate the pathogenic mechanism of POI. We created the W690C single-point mutation in the conserved TLDc domain of *C. elegans* LMD-3, which perfectly corresponds to a pathogenic NOCA7^W804C^ mutation found in human patients with POI^19^. We demonstrate that this single-point mutation significantly impairs reproductive capacity, causing reduced germline cell numbers and increased apoptosis. Mechanistically, our findings reveal that the mutation severely compromises LMD-3 protein stability and gene expression, leading to profound cellular autophagic-lysosomal dysfunction. This defect impairs both lysosomal acidification and its degradative capacity, causing the widespread accumulation of misfolded proteins and aggregates in the germline.

Critically, we show that this defect results in a functional B12 deficiency that can be corrected by mecobalamin (meCbl) supplementation, thereby restoring proteostasis. These findings provide critical insights into the molecular basis of female reproductive longevity, establishing a valuable genetic model for studying POI and highlighting a potential therapeutic target for mitigating age-related reproductive decline.

## Results

### Genetic and structural evidence for a shared POI-associated W690C mutation in LMD-3

The *C. elegans* LMD-3 protein, which serves as the sole OXR1 family member in the nematode, possesses a domain architecture highly conserved with its human orthologs, OXR1 and NCOA7 (Fig. 1A). This architecture is defined by three highly conserved LysM, GRAM, and TLDc domains. Despite the low 26.46% overall amino acid sequence identity between LMD-3 and human NCOA7^24^, the key functional domains, including the TLDc domain and other conserved motifs, are remarkably homologous (Supplementary Fig. 1).

**Fig 1.**
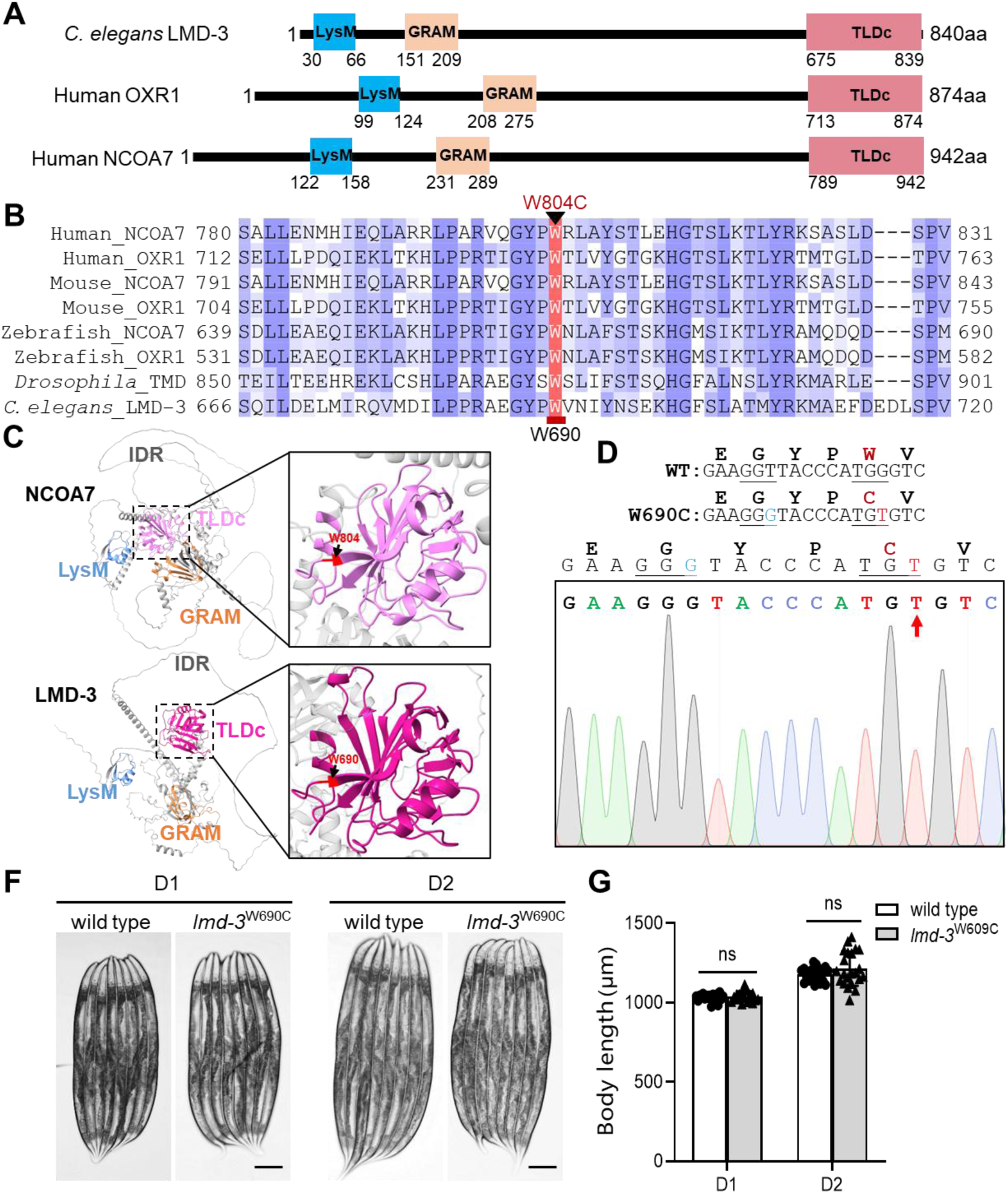
Characterization of the POI-associated W690C Mutation in the conserved TLDc domain of *C. elegans* LMD-3. **A** Schematic representation of conserved domains in *C. elegans* LMD-3 and its human orthologs, OXR1 and NCOA7. The schematic diagrams illustrate the conserved LysM, GRAM, and TLDc domains. The total number of amino acids (aa) is indicated for each protein. **B** The conserved tryptophan residue (W804) associated with premature ovarian insufficiency (POI) is highly conserved across species. A multiple sequence alignment of NCOA7 orthologs from five different species demonstrates the conservation of POI-associated W804 residue in human NCOA7. The residue, corresponding to W690 in *C. elegans* LMD-3, is highlighted with a red arrowhead. **C** Comparative structural analysis of the TLDc domain from human NCOA7 and *C. elegans* LMD-3. The TLDc domains are shown in detail to highlight their conserved structure similarity. The intrinsically disordered region (IDR) is colored gray. The conserved POI-associated tryptophan residues (human W804 and *C. elegans* W690) are indicated by black arrows and are colored ^W690C^ red in the detailed view. **D** Sequencing confirmation of the *C. elegans lmd-3* knock-in allele. A representative DNA sequencing trace shows the successful creation of the POI-associated W690C mutation. A red arrow marks the target base change. The target knock-in site is indicated by a red arrow. The resulting amino acid change (W to C) is highlighted in red, while a silent mutation introduced for screening is highlighted in light blue. **F**, **G** Development defects detection between wild-type and ^W690C^ *lmd-3* animals at D1 and D2 adult stages with bright-field image (**F**) and quantification (**G**) of animal body lengths (n ≥ 22 for each group, unpaired t-tests: ns, no significant difference). Scale bar, 50 μm.

The conservation of these domains is further highlighted by the POI-associated tryptophan residue (W804) in the TLDc domain of human NCOA7^19^. This residue is perfectly conserved across multiple species, including the corresponding W690 residue in *C. elegans* LMD-3 (Fig. 1B and Supplementary Fig. 1)^25^. The remarkable conservation of this clinically relevant residue across species provides a strong genetic basis for a shared pathogenic mechanism and validates the use of *C. elegans* W690C mutant as a model for human POI.

Following strong genetic evidence for a shared mechanism, we next sought to determine if this functional similarity was underpinned by structural conservation. A comparative structural analysis using AlphaFold3 revealed a remarkable similarity between the predicted *C. elegans* LMD-3 structure and its human NCOA7 ortholog (Fig. 1C). Despite differences in unique α-helices and intrinsic disordered regions (IDRs), the core LysM, GRAM, and TLDc domains were highly conserved. This structural homology was particularly striking in the V-ATPase-interacting TLDc domain, which showed an almost identical conformation and the conserved W804/W690 site (Fig. 1C and Supplementary Movie 1). These results provide compelling structural evidence that the TLDc-V-ATPase interaction is a central, conserved mechanism governing lysosomal activity and proteostasis across species.

Based on this strong evidence for a shared mechanism, we created an in vivo model of human POI by introducing the orthologous W690C mutation into *C. elegans lmd-3* using CRISPR/Cas9 (Fig. 1D and supplementary Table 1). A preliminary analysis of the *lmd-3*^W690C^ homozygous mutant animals revealed no obvious developmental defects, as confirmed by a lack of significant difference in body length compared to wild-type animals (Fig. 1F, G). The lack of significant developmental defects, consistent with the primary reproductive phenotype observed in human POI^19^, establishes the *lmd-3*^W690C^ strain as a high-fidelity in vivo model for investigating the molecular mechanisms of this disease.

### The POI-associated W690C mutation impairs fecundity and germline integrity in *C. elegans*

The deleterious W804C mutation in human NCOA7 has been shown to contribute to POI by accelerating oxidative stress-induced cellular senescence in ovarian granulosa cells^19^. Given the perfect conservation of this residue in *C. elegans* LMD-3 and the established role of this protein in oxidative stress resistance^20,26^, we investigated whether the orthologous *lmd-3*^W690C^ mutation would impair reproductive capacity in a manner consistent with the human disease. Our results showed a pronounced defect in fecundity, with the mutant strain exhibiting a 32.1% reduction in total progeny over their lifespan compared to wild-type animals (Fig. 2A and Supplementary Fig. 2A, B). Mirroring the rapid decline in fertility observed in human POI, this reduction was particularly striking during the peak reproductive period, suggesting that the mutation causes an accelerated and severe decline in fertility.

**Fig 2.**
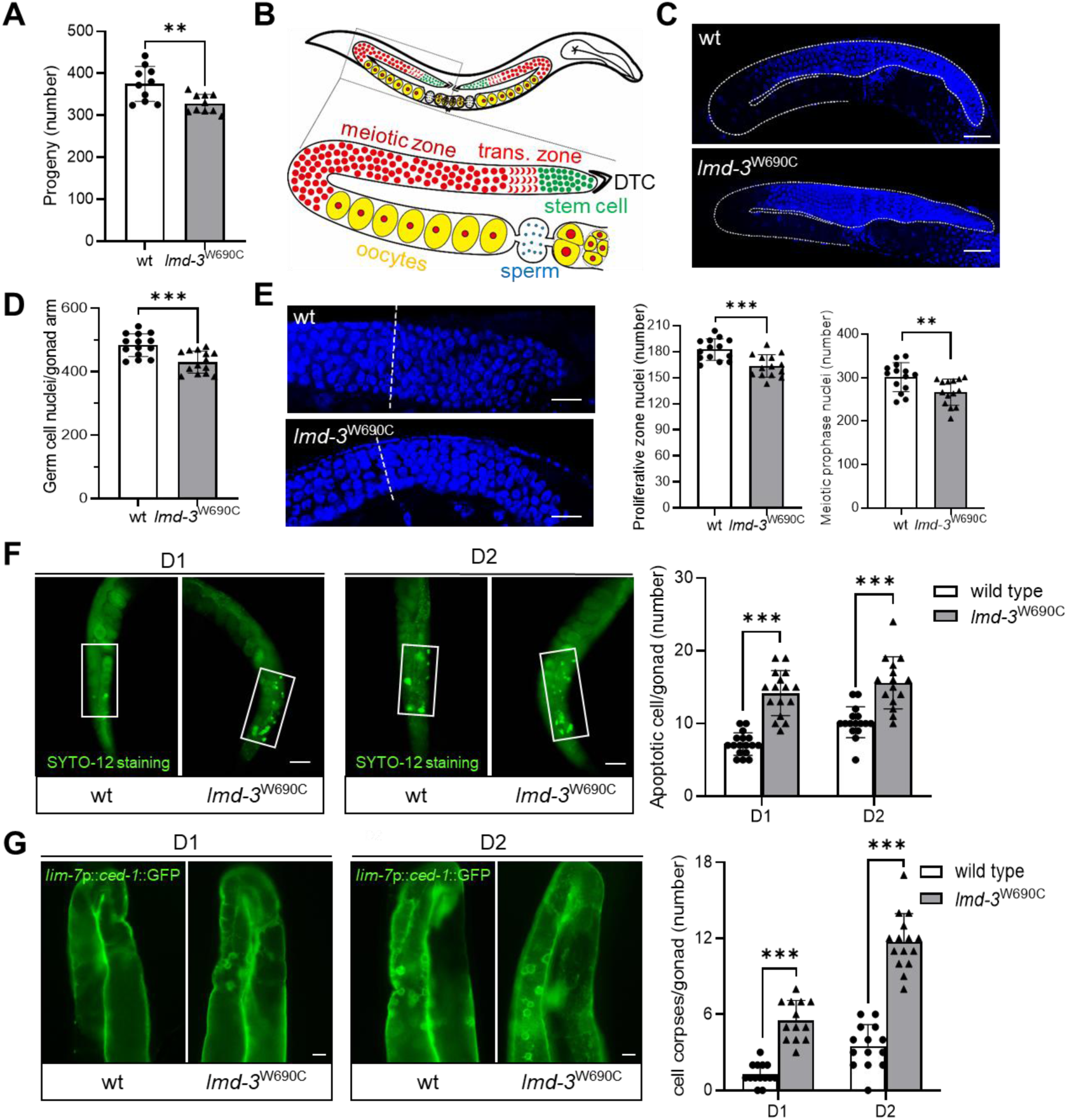
The POI-associated W690C mutation in *C. elegans* LMD-3 impairs fecundity and germline integrity. **A** Total brood sizes indicate the overall *^W690C^* reproductive ability of wild-type and *lmd-3* mutants. Data are presented as mean ± SEM (n = 10 per group, \*\**p* < 0.01). **B** Spatial organization of the C. elegans germline stem cell and meiotic regions. The schematic diagram illustrates one of the two distinct gonad arms, showing the spatial organization of the stem cell region, transition zone, and the meiotic phase along the distal-proximal axis. The distal tip cell (DTC) is also depicted, highlighting its role in enclosing the distal arm. **C, D** Representative fluorescence images (**C**) and quantification (**D**) of DAPI-stained total germline cells in wild-type and *lmd-3* mutants (n = 14 for each group, \*\*\**p* < 0.001). Scale bars, 50 μm. **D** Representative fluorescence images and quantification of DAPI staining for germline stem cells and meiotic cells in wild-type and *lmd-3* mutants (n = 14 for each group, \*\**p* < 0.01, \*\*\**p* < 0.001). Scale bars, 20 μm. **F, G** Germline apoptosis in wild-type and *lmd-3* mutants with SYTO12 staining (**F**) and a *lim-7p::ced-1::GFP* reporter (**G**). White arrowheads indicate apoptotic cell corpses. n ≥ 13 per group; unpaired t-tests, \*\*\**p* < 0.001. D1, Day 1 of adulthood; D2, Day 2 of adulthood. Scale bars, 50 µm.

To understand the cellular basis for this reduced fecundity, we next assessed germline integrity, hypothesizing that the pronounced reduction in fecundity was a direct result of germ cell deficiency^27^. The *C. elegans* germline is a dynamic, spatially organized tissue that produces all progeny. It is divided into distinct zones of proliferation, meiosis, and differentiation, all of which are essential for producing viable offspring (Fig. 2B). Germ cells are maintained by a distal stem cell region, regulated by the distal tip cell (DTC), which allows for clear visualization and analysis of germ cell progression throughout the gonad^28^.

Quantitative analysis of DAPI-stained gonads confirmed a significant reduction in the total number of germ cells in Day 1 adult *lmd-3*^W690C^ mutants (Fig. 2C, D). This defect was accompanied by a specific decline in the number of proliferative germline stem cells (Fig. 2E), indicating that the *lmd-3*^W690C^ mutation fundamentally compromises the maintenance of the germline progenitor pool, which is essential for ongoing germ cell production and homeostasis^29^.

To investigate the mechanism driving germ cell loss, we assessed germline apoptosis, a process frequently linked to oxidative stress^30^. Using two complementary methods, the nucleic acid dye SYTO12 and a *lim-7*p::*ced-1*::GFP reporter^31,32^, we observed a significant increase in apoptotic cell corpses within the germline of W690C mutants at Day1 and Day2 of adulthood (Fig. 2F, G and Supplementary Fig. 2C, D). The finding directly links increased germ cell death to reduced fecundity in this POI model. Our results collectively demonstrate the *lmd-3*^W690C^ mutation compromises germline integrity and reproductive capacity by promoting apoptosis, validating the mutant as a valuable in vivo model for human POI.

### The LMD-3 W690C mutation induces widespread and chronic cellular stress

To gain a deeper understanding of the physiological role of LMD-3, we first characterized its expression and subcellular localization. An integrated (Is) transcriptional reporter revealed that *lmd-3* is broadly expressed, including the nervous system, intestine, and vulva (Fig. 3A)^26^. Subcellular fractionation of the N-terminal mCherry-tagged LMD-3 protein and its TLDc domain confirmed their predominant localization to the cytoplasm and membrane fractions (Fig. 3B), consistent with its known association with lysosomes and the V-ATPase complex^22,23^. A minor nuclear fraction was also observed, which aligns with the transcriptional and antioxidant functions of its orthologs NCOA7 and OXR1 (Fig. 3B)^33,34^.

**Fig 3.**
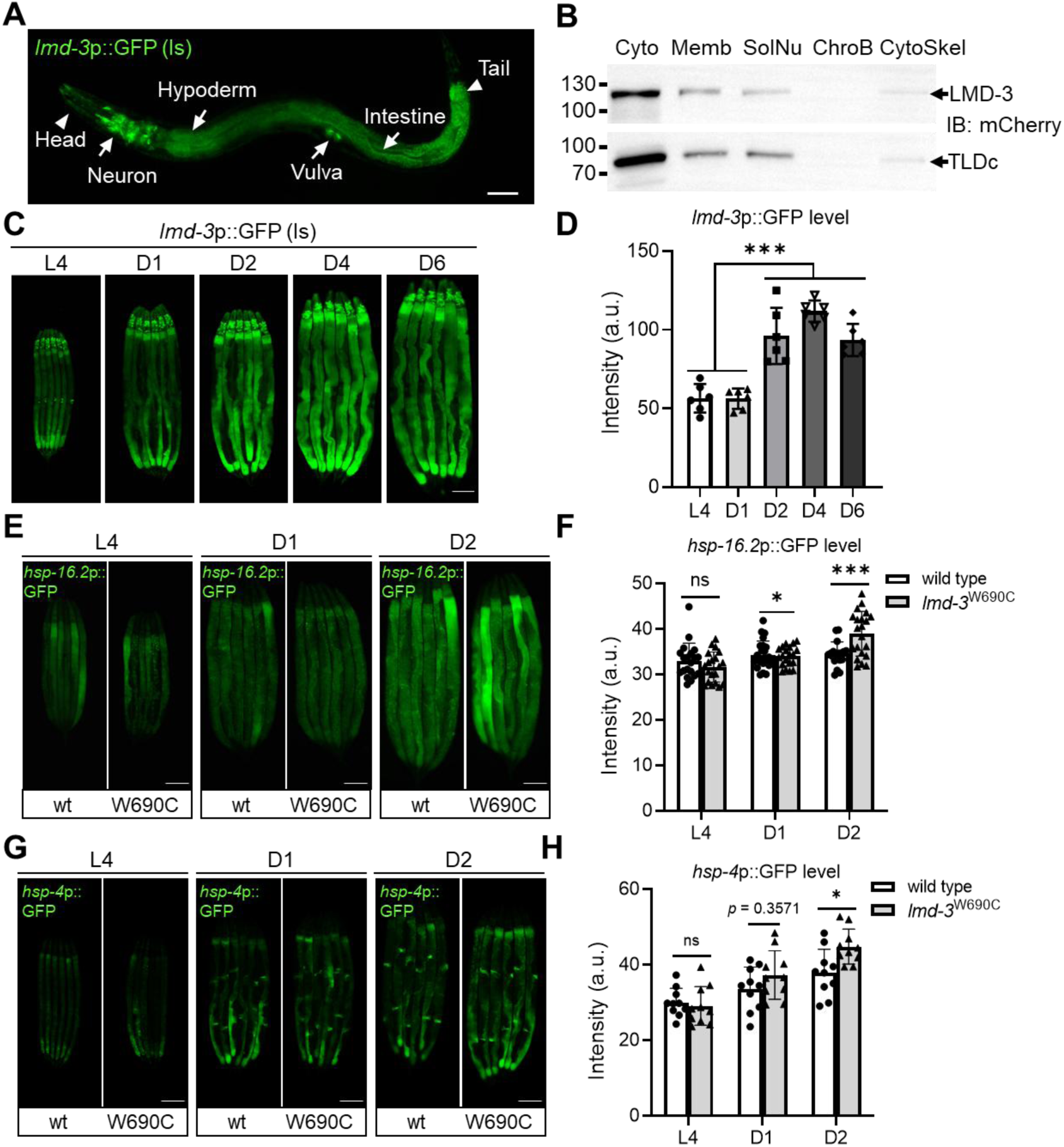
The expression pattern of LMD-3 and the role of W690C mutation in stress response. **A** Expression pattern of *lmd-3* at the L4 larval stage. Representative images show the expression of the *lmd-3p*::GFP transcriptional reporter). Is, integrated transgene. Scale bars, 100 µm. **B** Subcellular localization of full-length LMD-3 and its TLDc domain. The distribution of mCherry-tagged full-length LMD-3 (upper) and mCherry-tagged TLDc domain (lower) was analyzed. The fractions analyzed were cytoplasmic (Cyto), membrane (Memb), soluble nuclear (SolNu), chromatin-bound (ChroB), and cytoskeletal (CytoSkel). **C, D** Transcriptional level of *lmd-3* during different developmental stages with representative fluorescence images (**C**) and quantification (**D**) using the *lmd-3*p::GFP reporters (n ≥ 6 for each group, \*\*\**p* < 0.001). Is, integrated transgene. Scale bars, 100 μm. a.u., arbitrary units. **E, F** Representative fluorescence images (**E**) and quantification (**F**) of the UPR reporter *hsp-16.2*p::GFP in wild type and *lmd-3* animals at indicated stages (n ≥ 20 for each group, ns, no significant difference). Scale bars represent 100 μm. a.u., arbitrary units. L4, the fourth larval stage. D1, Day 1 of adulthood. D2, Day 2 of adulthood. **G, H** Representative fluorescence images (**G**) and quantification (**H**) of the UPR^ER^ reporter *hsp-4*p::GFP in wild-type and *lmd-3*^W690C^ animals at indicated stages (n ≥ 10 for each group, unpaired t-tests: ns, no significant difference, \**p* < 0.05). Scale bars represent 100 μm. a.u., arbitrary units.

Based on the established link between oxidative stress-induced cellular senescence and POI^19^, and given that the LMD-3 orthologs in yeast and humans are known to be stress-inducible^26,35^, we hypothesized that LMD-3 expression would increase with age as a protective response. Indeed, we found a gradual, age-dependent increase in *lmd-3* transcription (Fig. 3C, D), a pattern suggesting that LMD-3 is actively involved in preserving cellular health and longevity as the organism ages. This finding further supports a conserved role for LMD-3 as a protective factor against age-related stress, thereby promoting survival.

To determine if the W690C mutation compromises cellular stress resilience, we monitored cytoplasmic and endoplasmic reticulum (ER) stress throughout the late larval and early adult stages. We found that the cytosolic unfolded protein response (UPR^cyto^), monitored by the *hsp-16.2*p::GFP reporter^36^, showed a gradual but dramatic induction in the mutants (Fig. 3E, F and Supplementary Table 2), indicating an increase in oxidative stress. In parallel, the ER unfolded protein response (UPR^ER^), as indicated by the increased expression of the *hsp-4*p::GFP reporter^37^, also showed a progressive but apparent activation (Fig. 3G, H and Supplementary Table 2).

These results show that the POI-associated W690C mutation induces widespread but relatively mild and chronic cellular stress, as evidenced by the activation of both cytosolic and ER stress pathways. However, this general stress response is not sufficient to fully account for the early and severe reproductive decline observed in the mutant. This discrepancy suggests that the reproductive failure is not a secondary consequence of this general chronic stress but rather stems from a direct and severe breakdown in essential germline protein quality control, specifically through impaired lysosomal function and protein degradation^6,19,38^.

### The W690C-induced lysosomal dysfunction drives proteostasis collapse

Building on the established interaction between the TLDc domain and V-ATPases^22,23^, and the central role of lysosomes in aging, we investigated the impact of the W690C mutation on lysosomal function. Our transcriptional analysis revealed a significant downregulation of a wide range of lysosome-related genes in the mutant (Fig. 4A and Supplementary Fig. 3A). This included key components for acidification (V-ATPases), lysosomal structural integrity (LMP-2, NCR-1), and degradation (cathepsin proteases ASP-3, ASP-4). The initial finding suggests that the W690C mutation severely impairs the biogenesis and overall function of the lysosomal system.

**Fig 4.**
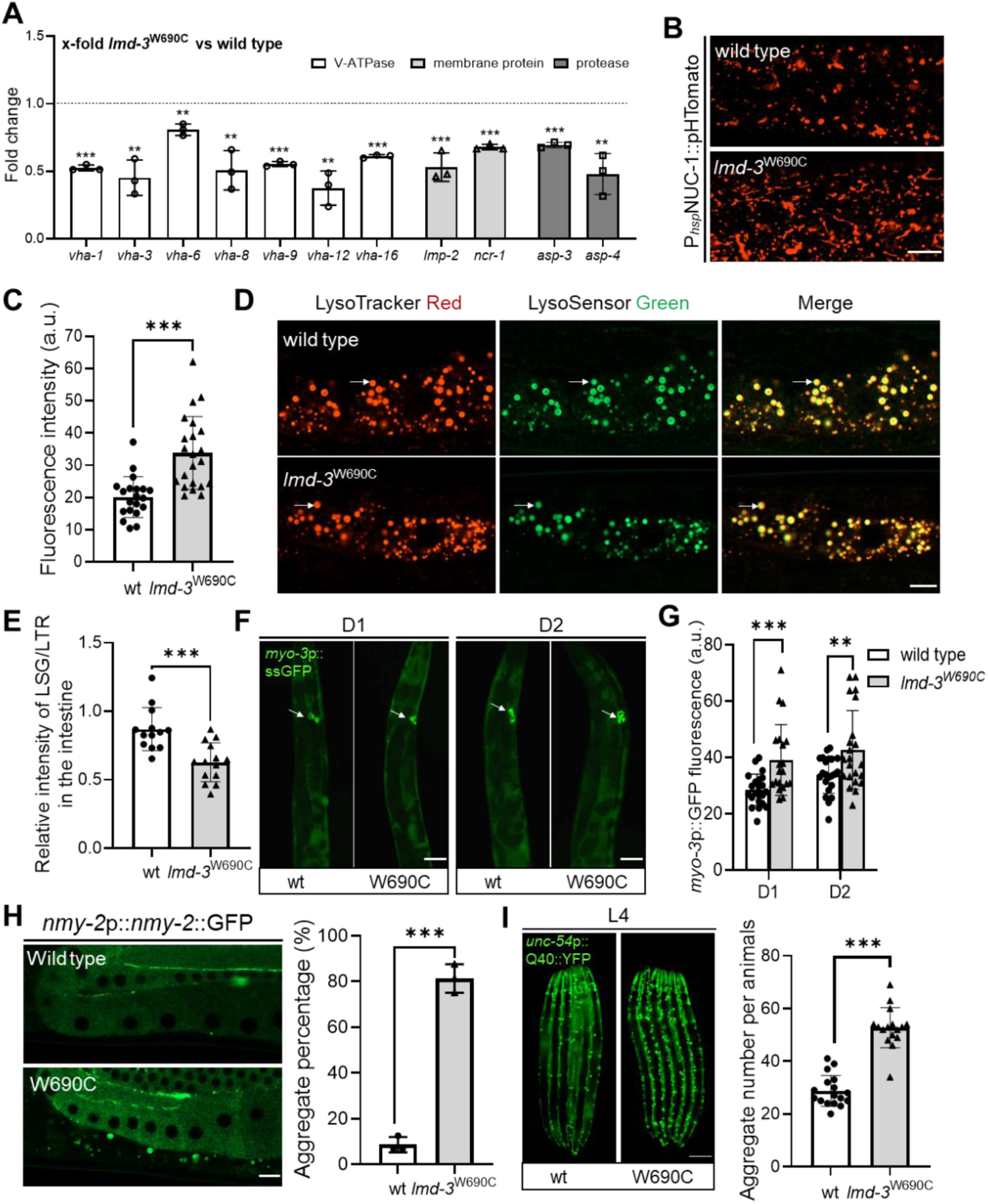
The W690C mutation affects proteostasis and lysosomal function in *C. elegans*. **A** Transcriptional analysis of lysosome-related genes in wild type and *lmd-3*^W690C^ animals at the D1 adult stage (n = 3 for each group, unpaired t-tests: \*\**p* < 0.01, \*\*\**p* < 0.001). **B, C** Representative fluorescence images (**B**) and quantification (**C**) of NUC-1::pHTomato expression in the hypodermis driven by a heat-shock (hs) promoter (n ≥ 20 per group, \*\*\**p* < 0.001). a.u., arbitrary units. Scale bar, 10 μm. **D, E** Representative confocal fluorescence images (**D**) and quantification (**E**) of intestines co-stained with Lysosome-Specific Green (LSG DND-189) and Lysosome-Targeting Red (LTR DND-99) in wild type and *lmd-3* animals at the D1 adult stage. White arrowheads indicate vesicular lysosomes stained by both dyes. Relative intensity of LSG/LTR were quantified (n ≥ 13 per group, \*\*\**p* < 0.001). Scale bars, 10 μm. **F, G** Representative fluorescence images (**F**) and quantification (**G**) of secreted protein accumulation (*myo-3*p::ssGFP) in the coelomocytes of wild type and *lmd-3* animals at the D1and D2 stages (n ≥ 18 per group, unpaired t-tests: \*\**p* < 0.01, \*\*\**p* < 0.001). Scale bars, 50 μm. a.u., arbitrary units. **H** Confocal fluorescence images and quantification of NMY-2::GFP accumulation in wild type and *lmd-3* animals at D1 stage (mean ± SEM, n = 3 biological replicates, 10 animals per condition per replicate, unpaired t-tests: \*\*\**p* < 0.001). **I** Fluorescence images and quantification of Q40::YFP accumulation in the body wall muscles in wild type and *lmd-3*^W690C^ animals at the L4stage (n ≥ 16 per group, unpaired t-tests: \*\*\**p* < 0.001). Scale bars, 50 μm.

To directly confirm that the transcriptional defects led to a loss of lysosomal function, we measured lysosomal acidification, a process critical for protein degradation^39^. The use of a heat-shock-inducible pH-sensitive reporter, NUC-1::pHTomato^40^, revealed a significantly higher fluorescence intensity in the mutant, indicating a less acidic, more neutralized lysosomal pH (Fig. 4B, C and Supplementary Fig. 3B, C). This defect was independently corroborated by co-staining with the acidotropic dye LysoSensor Green (LSG) and the lysosome-specific dye LysoTracker Red (LTR)^41^. The resulting reduced LSG/LTR fluorescence ratio in the mutant definitively confirmed that the transcriptional defects resulted in a severe impairment of lysosomal acidity in *C. elegans* (Fig. 4D, E and Supplementary Fig. 3D, E).

The consequence of the acidification defect was a severe breakdown in degradative capacity, which we assessed by monitoring the clearance of secreted ssGFP^42^. The reporter showed a significant accumulation of fluorescence in the coelomocytes of the mutant (Fig. 4F, G), demonstrating a failure of lysosomal degradation. This defective degradation led to a broad collapse in proteostasis, as evidenced by extensive aggregation of the age-related protein NMY-2::GFP in oocytes^43^ (Fig. 4H) and the polyglutamine reporter Q40::YFP in body wall muscles^44^ (Fig. 4I). The vital role of LMD-3 in protein homeostasis was further highlighted by the accumulation of vitellogenin, a major egg yolk protein, in the mutant’s intestine, pseudocoelom, and germline (Supplementary Fig. 3F), confirming that LMD-3 deficiency impairs the processing of this crucial reproductive protein.

To establish that the lysosomal defect is a direct cause of the reproductive failure, we used the *vha-12(ok821)* mutant to genetically neutralize V-ATPase activity. This loss-of-function allele not only recapitulated the reproductive defects observed in the W690C mutant but also exacerbated the decline (Supplementary Fig. 3G). These findings cement a causal link between the W690C-induced V-ATPase dysfunction and reproductive decline, underscoring the vital role of LMD-3 in maintaining lysosomal function for reproductive longevity.

### Vitamin B12 restores proteostasis by repairing LMD-3(W690C)-mediated autophagic-lysosomal defects

Following our discovery of profound lysosomal dysfunction and proteostasis collapse in the LMD-3(W690C) mutant, which mirrors the defective autophagy and senescence linked to NCOA7 dysfunction in human POI^19^, we investigated the integrity of the autophagic pathway. We first utilized a transcriptional reporter for the stress-inducible autophagy gene, *tts-1*^45^. In contrast to wild-type animals, the W690C mutants showed a significant reduction in *tts-1*p::GFP fluorescence (Fig. 5A). This was further supported by the notable downregulation of key autophagy genes, including the master transcription factor *hlh-30*^46^, along with genes essential for autophagosome formation, cargo recognition, fusion, and lysosomal degradation (Fig. 5B, Supplementary Fig. 5A). These findings suggested that the W690C mutation severely compromises the biogenesis and overall function of the autophagic system.

**Fig 5.**
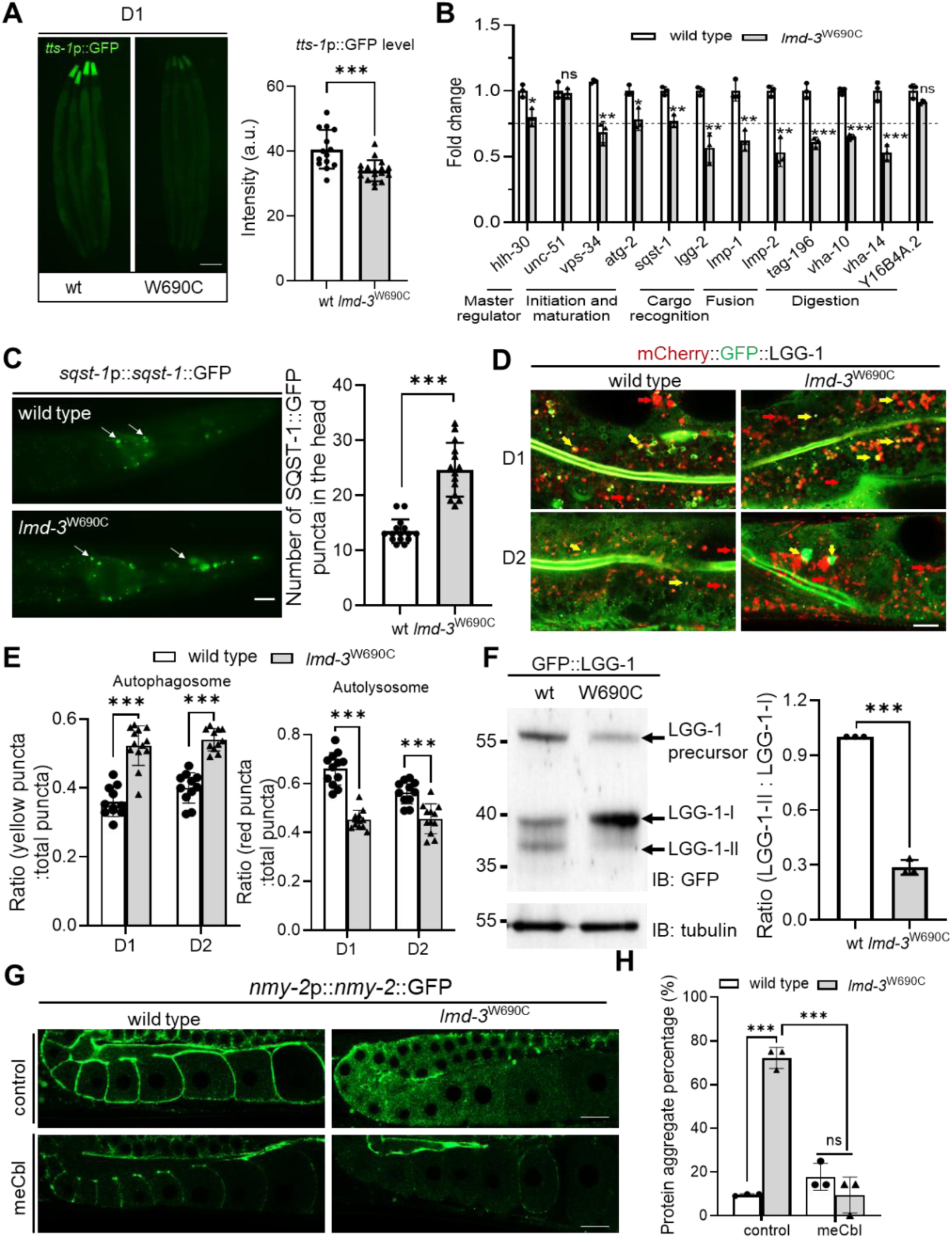
meCbl restores proteostasis in LMD-3(W690C) mutants by repairing autophagic-lysosomal defects. **A** Representative fluorescence images and quantification of the autophagy reporter *tts-1*p::GFP in wild type and *lmd-3* animals at the indicated stages (n ≥ 10 per group, \*\*\**p* < 0.001). a.u., arbitrary units. Scale bars represent 100 μm. **B** Transcriptional analysis of key autophagy genes was performed in wild type and *lmd-3* animals, and gene expression was quantified by RT-qPCR (n = 3 per group, unpaired t-tests: \**p* < 0.05, \*\**p* < 0.01, \*\*\**p* < 0.001, ns, no significant difference). **C** Representative fluorescence images and quantification of SQST-1::GFP accumulation in wild type and *lmd-3* animals at D1 adult stage (n ≥ 14 per group, \*\*\**p* < 0.001). White arrowheads denote fluorescent protein aggregates. Scale bars, 50 μm. **D, E** Representative confocal images (**D**) and quantification (**E**) of the mCherry::GFP::LGG-1 reporter in wild type and *lmd-3* animals at D1 and D2 adult stages. The ratio of autophagosomes (yellow puncta, yellow arrow) to total puncta and autolysosomes (red puncta, red arrow) to total puncta was quantified (n ≥ 11 per group, \*\*\**p* < 0.001). Scale bar, 10 μm. **F** Representative Western blot and corresponding quantification of the ratio of lipidated LGG-1-II to unlipidated LGG-1-I protein in wild-type and *lmd-3* mutant animals (n = 3 biological replicates, \*\*\**p* < 0.001). a.u., arbitrary units. IB, immunoblotting. **G, H** Confocal fluorescence images (**G**) and corresponding quantification (**H**) of *nmy-2*p::*nmy-2*::GFP accumulation in the oocytes*^W690C^* of wild-type and *lmd-3* mutants at D1 of adulthood, with mecobalamin (meCbl) supplementation (mean ± SEM, n=3 biological replicates, animals per condition per replicate = 10, \*\*\**p* < 0.001, ns, no significant differences). Scale bars, 10 μm.

To confirm that this transcriptional defect leads to a functional impairment, we proceeded to examine the accumulation of SQST-1, the *C. elegans* ortholog of p62. Efficient clearance of SQST-1 is indicative of a healthy autophagic flux, and we observed a massive accumulation of SQST-1::GFP aggregates in the adult W690C mutants (Fig. 5C and Supplementary Fig. 4B), providing strong evidence that the overall autophagic degradation pathway is severely compromised.

To precisely characterize autophagic failure, we first utilized tandem fluorescent reporter mCherry::GFP::LGG-1 to distinguish between autophagosomes formation (yellow puncta) and maturation into autolysosomes (red puncta), where acid-sensitive GFP is quenched in the acidic lysosomal environment^47^. Confocal analysis revealed a dramatic increase in yellow puncta and a parallel decrease in red puncta in the W690C mutant (Fig. 5D, E and Supplementary Fig. 4C, D). This block in autophagic flux at the fusion/degradation stage directly reinforces the severe lysosomal acidification defect observed previously (Fig. 4B-E). Furthermore, GFP::LGG-1 Western blot analysis revealed a simultaneous defect in autophagosome biogenesis^48^, shown by the significant drop in the ratio of membrane-associated LGG-1-II (lipidated) to cytosolic LGG-1-I (unlipidated) (Fig. 5F). These findings confirm the W690C mutation induces a comprehensive, multi-stage failure of the autophagic-lysosomal pathway.

Given that LMD-3^W690C^ dysfunction impairs lysosomal function and reproductive health, we hypothesized that enhancing cellular resilience could restore proper lysosomal function. Since lysosomes are critical for the uptake and processing of dietary vitamin B12, the catastrophic lysosomal failure in *lmd-3* mutants would result in a state of functional B12 deficiency^49^. We therefore tested the capacity of the active form, meCbl, to mitigate this lysosomal dysfunction. meCbl supplementation significantly rescued the accumulation of the aggregation-prone protein NMY-2::GFP in the germline (Fig. 5G, H) and improved protein clearance in coelomocytes, as evidenced by reduced ssGFP fluorescence (Supplementary Fig. 4E, F). Collectively, these results demonstrate that B12 supplementation mitigates cellular stresses and directly improves the defective lysosomal function, thereby restoring germline proteostasis and protein clearance in *lmd-3*^W690C^ mutants.

### The W690C mutation compromises LMD-3 structure and expression

Having demonstrated the essential role of the W690C mutation in disrupting lysosomal function and cellular homeostasis in *C. elegans*, we set out to determine the precise molecular basis of this effect. We first investigated whether the mutation affected LMD-3 expression at the transcriptional level. Using primers designed to target three distinct locations along the gene (Fig. 6A and supplementary Table 3), we detected a significant and consistent reduction in the overall *lmd-3* transcript level in the mutant strain compared to the wild type (Fig. 6B and Supplementary Fig. 5A). This reduction in transcript levels exacerbates the loss-of-function phenotype.

**Fig 6.**
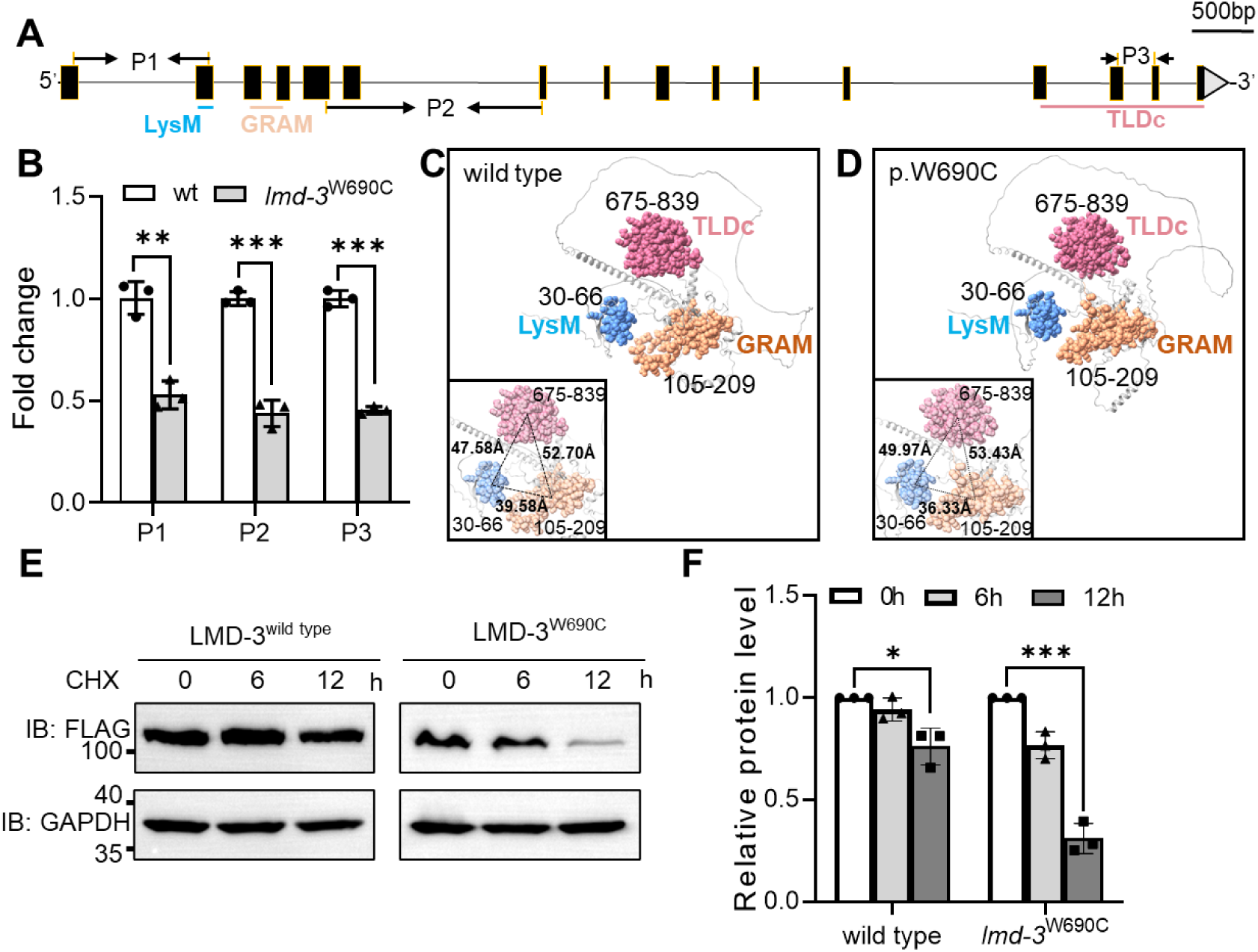
The W690C mutation affects LMD-3 protein structure and stability. **A** The schematic diagram shows the genomic structure of *lmd-3*, including the positions of its conserved LysM, GRAM, and TLDc domains. The locations of the qRT-PCR primers (P1, P2, P3) designed to quantify lmd-3 mRNA are also shown. **B** qRT-PCR measurements for endogenous *lmd-3* mRNA levels in wild type (wt) and *lmd-3* animals (n = 3 per group, unpaired t-tests: \*\**p* < 0.01, \*\*\**p* < 0.001)**. C, D** Structural analysis of the LMD-3 protein. Predicted three-dimensional structures of wild-type (**C**) and W690C (**D**) proteins are presented to illustrate the effects of the W690C mutation on protein folding. Key domains are color-coded: LysM (cornflower blue), GRAM (light salmon), and TLDc (pale violet red). A magnified view shows the trimer-like region, where the mutation leads to altered inter-subunit distances in the p.W690C protein (49.97 Å, 53.43 Å, 36.33 Å) compared to the wild-type (47.58 Å, 52.70 Å, 39.58 Å). **E, F** The W690C mutation reduces LMD-3 protein stability. Western blot images (**E**) and semi-quantitative analysis (**F**) show the turnover of 1xFLAG-tagged LMD-3 in H293T cells treated with cycloheximide (CHX) for 0, 6, and 12 h (n = 3 biological replicates, \**p* < 0.5, \*\*\**p* < 0.001). a.u., arbitrary units.

With the established link between human POI-associated NCOA7 mutations and protein stability^19^, we reasoned that the W690C mutation would compromise the structure of LMD-3. To confirm this hypothesis, three-dimensional structural models of the *C. elegans* wild-type and W690C-mutated LMD-3 protein were developed^50^. Our model showed that the LMD-3 protein, like its human ortholog NCOA7, forms a conserved trimer-like region comprised of its LysM, GRAM, and TLDc domains (Fig. 6C). Although the overall core domains maintained their general conformation, we found that the W690C mutation significantly altered the inter-subunit distances within the predicted trimer-like region (Fig. 6D). This structural change suggests a destabilization of a key binding interface. Critically, six machine learning-based methods confirmed that the W690C mutation leads to a compromise in overall protein stability (supplementary Table 4)^51–55^, consistent with findings from a human study^19^.

To directly confirm that the W690C mutation results in reduced protein stability, we performed a cycloheximide (CHX) chase assay to evaluate the turnover of LMD-3 protein^56^. As a result, while wild-type LMD-3 protein remained relatively stable, the mutant protein exhibited a significant decrease in levels, showing a marked degradation after 12 hours of CHX treatment (Fig. 6E, F and Supplementary Fig. 5B). This result provides compelling evidence that the W690C mutation compromises the stability of LMD-3 protein, mechanistically linking the structural defect to its rapid degradation. Collectively, these findings establish that the POI-associated W690C mutation compromises both *lmd-3* gene expression and protein stability, thereby leading to the lysosomal defects and reproductive decline.

## Discussion

Reproductive longevity is a fundamental aspect of healthy aging, yet its molecular mechanisms remain incompletely understood. Our study establishes a novel POI-associated *C. elegans* model to dissect the pathophysiological basis underlying fertility decline. By introducing an orthologous W690C mutation into LMD-3, the single-gene family member of human NCOA7 and OXR1, we found compelling evidence that this genetic lesion impairs reproductive capacity by compromising protein stability and inducing a profound collapse in cellular proteostasis. These findings define a specific pathogenic mechanism in which a single amino acid substitution leads to widespread cellular dysfunction, thereby directly linking a conserved protein to the maintenance of reproductive health.

Our study reveals that the W690C mutation, located within the conserved TLDc domain, severely compromises LMD-3 protein stability. This finding provides a crucial mechanistic link between the genetic mutation and the observed loss-of-function phenotypes. A key insight from our data is that this mutation also severely reduces the overall *lmd-3* mRNA level (Fig. 6B), which directly contributes to the loss-of-function phenotype. This observation contrasts sharply with the human NCOA7^W804C^ mutation, where mRNA levels are unaffected^19^. This observed discrepancy in mRNA levels is a critical point that highlights the mechanistic advantage of studying a non-redundant system. In mammals, the presence of the ortholog OXR1 likely provides cellular buffering or compensation that masks any regulatory feedback related to NCOA7 loss. As *lmd-3* is the sole member of this gene family in *C. elegans*, its dysfunction has a more direct, non-redundant, and severe impact, making our model an ideal system to study the core pathogenic mechanism.

The destabilization of LMD-3^W690C^ protein leads to profound lysosomal dysfunction, characterized by defective acidification and impaired degradative capacity. These data align with emerging evidence that lysosomes are not merely “waste disposal plants” but critical signaling hubs whose dysfunction is linked to aging and neurodegenerative diseases^57–59^. Our study now unequivocally links organelle failure to a major defect in reproductive capacity, thereby establishing a direct causal link. This primary lysosomal defect triggers a downstream cascade of cellular damage, as the accumulation of misfolded proteins and aggregates creates a proteotoxic environment that is detrimental to sensitive, non-dividing cells like oocytes^6,60^. Such chronic cellular stress eventually manifests as increased germ cell apoptosis and a reduction in total cell number, fully accounting for the decline in reproductive capacity in the mutant. Ultimately, our findings reveal an elegant pathway linking impaired proteostasis, a well-known hallmark of aging^61^, to the specific context of ovarian aging.

Beyond identifying the pathogenic mechanism, our study provides pioneering evidence for a therapeutic strategy targeting this conserved pathway. Given that lysosomal failure compromises the cellular uptake and processing of vitamin B12, the LMD-3^W690C^ mutation creates a state of functional B12 deficiency that subsequently disrupts propionate metabolism. This is critical because B12 is an essential cofactor for the enzyme methylmalonyl-CoA mutase, which catalyzes a required step in propionate metabolism^62^. A functional deficiency in meCbl impairs this pathway, which is known to lead to toxic metabolite accumulation and cellular stress, thereby exacerbating the lysosomal dysfunction^63^. Supplementation with the active form, meCbl, successfully rescued the LMD-3^W690C^-induced proteostasis collapse, directly restoring germline protein homeostasis and enhancing clearance. This highly significant finding suggests a simple nutritional supplement could mitigate the effects of this severe genetic dysfunction, highlighting the immediate clinical relevance of our *C. elegans* model.

Our human disease-driven *C. elegans* model represents a significant advancement in the study of reproductive aging. By establishing a direct causal link between a specific genetic lesion, lysosomal dysfunction, proteotoxicity, and reproductive failure, our model moves beyond correlational studies. Furthermore, the model is highly relevant to human POI, as it recapitulates key features of the human condition, including germ cell depletion and premature reproductive decline^64^.

Beyond the specific W690C mutation, our work suggests that LMD-3/NCOA7 and its genetic interactors are promising candidates for genetic screening in patients with idiopathic POI. Furthermore, our findings raise broader questions about the role of wild-type LMD-3/NCOA7 in age-related fertility decline. Recent evidence has shown that the expression and function of this protein naturally wane with age, leading to a gradual decline in the lysosomal and proteostasis health observed in aged oocytes^19^. Consequently, our mutant model serves as an accelerated version of the natural aging process, providing a powerful system for dissecting these fundamental mechanisms.

The *C. elegans* model we established provides an invaluable high-throughput in vivo platform for screening potential drug interventions that can enhance lysosomal activity through mechanisms like TFEB activation or other protein homeostasis-restoring pathways like autophagy and proteostasis^47,65,66^. Furthermore, while our study establishes LMD-3 as a crucial regulator of lysosomal activity, its precise biochemical function within this organelle remains to be fully elucidated. Therefore, further studies are required to identify its binding partners and determine whether it acts as a scaffold, a regulator, or serves another function in maintaining lysosomal pH and enzymatic activity.

In summary our study establishes that the POI-associated W690C mutation in LMD-3, supported by genetic and structural evidence, compromises protein stability and expression, leading to a profound collapse in lysosomal function. This collapse triggers proteotoxic stress, germ cell death, and reproductive failure. Crucially, we define the resultant phenotype as a functional B12 deficiency, rationalizing mecobalamin supplementation as a viable therapeutic strategy. This discovery provides a valuable in vivo platform for screening potential therapeutic compounds. Furthermore, it establishes lysosomal activity as a crucial determinant of reproductive longevity and a promising target for extending reproductive capacity.

## Methods

### *C. elegans* strains and maintenance

We maintained all *C. elegans* strains (Supplementary Table 1), including the wild-type N2 Bristol strain and all other strains, at 20°C on standard nematode growth medium (NGM) plates. These plates were uniformly seeded with *Escherichia coli* OP50 as a food source. To ensure experimental synchrony, we obtained age-matched populations by treating gravid adults with a bleaching solution to isolate and hatch eggs.

### CRISPR-Cas9 mediated mutagenesis

The *lmd-3*^W690C^ point mutation was generated by SunyBiotech using the established CRISPR-Cas9 gene editing system. The presence of the desired point mutation was then definitively confirmed by Sanger sequencing (Supplementary Table 3). To remove any potential off-target mutations and ensure a clean genetic background, the verified mutant was subsequently outcrossed to the wild-type N2 strain three times before all phenotypic and functional analyses commenced.

### Plasmid construction

To facilitate biochemical analyses in mammalian cells, we generated expression vectors for 1xFLAG-tagged LMD-3 proteins. The full-length *lmd-3* cDNA sequence were amplified by PCR from both the wild-type and *lmd-3*^W690C^ mutant strains. These amplified fragments were subsequently subcloned into the pcDNA3.1 vector. Primers used for the amplification are listed (Supplementary Table 3).

### Quantitative real-time PCR (qRT-PCR)

To quantify target gene transcripts, 200 synchronized Day 1 adult animals were collected into M9 buffer, lysed using Trizol Reagent (Invitrogen, 15596026CN), and immediately frozen in liquid nitrogen. Total RNA was subsequently isolated via standard chloroform extraction and isopropanol precipitation. A total of 1.2 μg isolated RNA was then reverse-transcribed into complementary DNA (cDNA) using the HiScript III 1st Stand cDNA Synthesis Kit (+ gDNA wiper) (Vazyme, R312-01). Quantitative real-time PCR (qRT-PCR) was performed on an ABI QuantStudio 1 machine using ChamQ SYBR Color qPCR Master Mix (Vazyme, Q411-02). Transcript levels for the target genes were calculated and normalized to the endogenous control gene, *act−1*.

### Reproductive span analysis

As previously described, a minimum of 10 synchronized hermaphrodite larvae at L4 stage for each strain were individually transferred to fresh NGM plates at every 12 hours^67^. This transfer process continued until the cessation of reproduction, allowing for the isolation of all laid eggs. The number of progenies was counted at each transfer, and the total number of progenies per animal was calculated. The experiment was performed in three biological replicates.

### Measurement of body length

Synchronized *C. elegans* at designated developmental stages were first anesthetized using levamisole and mounted onto agarose pads. Images were captured using a Nexcope 910 microscopy system. The linear distance from the head to the tail of each worm was then measured using the Image View software to determine body length. A minimum of 22 worms were analyzed for each group.

### Microscopy and imaging analysis

For all standard fluorescence and bright-field microscopy experiments, synchronized animals were first anesthetized using levamisole and mounted onto agarose pads for imaging. Images were captured using either a Nexcope 910 fluorescence microscope or a high-resolution laser scanning confocal microscope (Carl Zeiss, LSM 980), depending on the required resolution and image type. Subsequently, fluorescence intensity and other quantitative metrics, such as puncta number or size, were measured and analyzed using the ImageJ software package.

### Germline staining and quantification

To analyze the integrity and morphology of the germline, synchronized Day 1 adult animals were first fixed by immersion in ice-cold methanol for 5 minutes. Following the removal of methanol, the samples were washed three times with M9 buffer. Germ cell nuclei were then stained by incubating the samples in a DAPI solution (Macklin, D807022) for 30 minutes in the dark, followed by three additional washes. The stained worms were then mounted, and their gonads were examined using a laser scanning confocal microscope. The total number of germ cells and the specific count of proliferative germline stem cells in each gonad were subsequently determined quantitatively using ImageJ software.

### SYTO12 staining for germline apoptotic cells

Synchronized larvae from control and experimental groups were collected and allowed to develop into Day 1 and Day 2 adults. These adult animals were then stained by incubation in SYTO 12 dye (Thermo Fisher Scientific, S7575) in the dark for 3 hours. Following staining, worms were washed three times with buffer via centrifugation (4,000 rpm, 1 minute) and transferred to fresh NGM plates for a 1-hour recovery period to clear dye-labeled bacteria from the intestines. Finally, the worms were anesthetized with levamisole, mounted on agarose pads, and imaged using an objective under green fluorescence microscopy to count the apoptotic cell corpses.

### Dual lysosomal staining and quantification

Synchronized Day 1 adult animals were co-stained for 1 hour at 20°C in the dark with 10 mM LysoSensor Green DND-189 (Yeasen) and LysoTracker Red DND-99 (Yeasen) as previously described^68^. The animals were transferred to fresh NGM plates seeded with OP50 for a 1-hour recovery period to clear residual dye from the gut bacteria. Lysosomal acidification was then quantitatively evaluated by measuring the relative fluorescence intensity ratio of LysoSensor Green to LysoTracker Red using ImageJ software, where a lower ratio indicates reduced lysosomal acidity.

### Cell culture and Transient transfection

HEK-293T cells were maintained in Dulbecco’s Modified Eagle’s Medium (DMEM) (Gibco, C119955-00BT) supplemented with 10% fetal bovine serum (FBS) (Excell, FSP-500) in a humidified atmosphere at 37°C with 5% CO₂. For transient transfection experiments, cells were transfected with the appropriate plasmid DNA using polyethyleneimine (PEI) (Sigma-Aldrich, 408727). The transfection mixture was prepared using a DNA-to-PEI mass ratio of 1:4.5, with a total of 1 µg DNA per 10^6 cells used for each condition.

### Cycloheximide pulse chase assay

The compromised stability of the 1xFLAG-tagged protein was evaluated in cells using a Cycloheximide (CHX) chase assay, which monitors protein turnover^69^. Thirty-six hours after transfection, cells were subjected to 50 µg/mL CHX to block protein synthesis. At designated time points, cells were collected, and the degradation of wild-type and mutant were determined by quantifying protein levels via Western blotting.

### Autophagy flux and aggregation analysis

To assess autophagy flux, synchronized worms expressing the tandem fluorescent reporter mCherry::GFP::LGG-1 were used. Puncta were counted in the anterior intestinal cells, where the number of autophagosomes (AP) was determined by counting the puncta positive for both the GFP and mCherry (yellow/orange), and the number of autolysosomes (AL) was determined by counting mCherry-only positive puncta (red, due to quenching in the acidic lysosome). Additionally, to evaluate the accumulation of aggregated cargo, the number of SQST-1::GFP positive puncta was counted specifically within the head neurons of synchronized worms. Both assays were conducted with a minimum of 11 worms examined per condition across three independent experiments.

### Mecobalamin supplementation

A 150 μM stock solution of meCbl (Macklin, M812748) was initially prepared in H₂O (which also served as the control solvent). This stock solution was then diluted to a final working concentration of 0.15 μM directly into the NGM agar prior to pouring the plates, ensuring the worms were exposed to the compound throughout their life cycle.

### Quantification of *myo-3*p::ssGFP intensity

Fluorescence images were captured from synchronized adult *C. elegans* at Day 1 of adulthood expressing the secreted reporter *myo-3*p::ssGFP using Nexcope microscopy. The average fluorescence intensity of the ssGFP signal was quantified using ImageJ software. At least 18 animals were analyzed for each strain and condition.

### Quantification of NUC-1::pHTomato intensity

Synchronized Day 1 adult *C. elegans* expressing P*_hs_*NUC-1::pHTomato were first subjected to a 33°C heat shock for 30 minutes to induce expression. Following induction, animals were allowed to recover at 20°C for 24 hours before imaging^40,43^. The average intensity of pHTomato was then quantified per area in the hypodermis using ImageJ software. A minimum of 20 worms were analyzed for each strain.

### Western blotting

A total of 200 synchronized day 1 adult worms were collected, lysed directly in 1× sample buffer, and boiled at 95°C for 10 minutes. The resulting protein extracts were separated by SDS-polyacrylamide gel electrophoresis (SDS-PAGE) and transferred onto a PVDF membrane. The membrane was blocked for 1 hour at room temperature using 5% non-fat milk in TBST before being incubated overnight at 4°C with the following primary antibodies: anti-GFP rabbit polyclonal antibody (1:1000, abclonal, AE011), anti-α-tubulin mouse monoclonal antibody (1:5000, Sigma-Aldrich, T5168), anti-flag DDDDK-tag mAb antibody (1:10000, MBL, M185) and anti-GAPDH antibody (1:500, Absin, abs830030ss). After washing, the membrane was incubated with HRP-conjugated secondary antibodies for 1 hour at room temperature. Finally, protein bands were visualized using an enhanced chemiluminescence (ECL) detection system.

### Protein Sequence and Structure Prediction

The protein sequences for human NCOA7 (NCBI accession: NP_001186548.1) and its *C. elegans* ortholog LMD-3 (NCBI accession: NP_505173.2) were retrieved from the National Center for Biotechnology Information (NCBI) database. Comparative three-dimensional (3D) structural modeling for wild-type and the W690C mutant was performed using. The predicted structures were visualized and analyzed using the ChimeraX 1.10.1 software package.

### Inter-domain distance measurement

To quantify the structural impact of the mutation on the predicted LMD-3 trimer-like region, the distances between the conserved, and domains were measured. The centroid of each respective domain was defined as a vertex of a triangle. The distances between these centroids were then calculated using the “Distance” function within the “Annotation” module of ChimeraX 1.10.1.

### Protein stability prediction

To assess the effect of the point mutation on protein stability, we employed six distinct machine learning-based prediction tools^51–55^. The LMD-3 protein sequence and 3D structural model were the same as those used for the structural analysis above. For each web-based tool, the W690C substitution was specified to calculate the change in free energy (ΔΔG). The definition of a “Destabilizing” prediction varied among the different websites, corresponding to specific ΔΔG thresholds.

### Statistical analysis

All data were statistically analyzed using GraphPad Prism 9. Data are presented as the mean, with error bars representing the standard error of the mean (SEM). Student’s t-tests were utilized for comparisons between two distinct groups, while one-way ANOVA tests were performed for multiple group comparisons. Statistical significance was established at *p* < 0.05 (indicated by *), *p* < 0.01 (indicated by **), and *p* < 0.001(indicated by ***).

## Data availability

All data supporting the findings of this study, presented within the main manuscript and the Supplementary Information, are available in the accompanying Source data file provided with this paper. All unique reagents and strains generated throughout this study are available from the corresponding author upon reasonable request and the completion of a standard Material Transfer Agreement (MTA).

## Acknowledgements

We thank the Caenorhabditis Genetics Center (CGC), Dr. Xiaochen Wang (Southern University of Science and Technology), Dr. Dengke Ma (UCSF), and Dr. Chenggang Zou (Yunnan University) for strains. Dr. Yingying Qin, Dr. Ting Dong, and Dr. Xue Jiao for their valuable suggestions (Shandong University).

## Author contributions

Conceptualization: W.L, X.W, Z.Z; Data Curation: Y.Z, T.W, Z.Z; Formal Analysis: Y.Z, T.W; Funding Acquisition: Z.Z; Investigation: Y.Z, T.W, M.G, W.L, X.W, Z.Z; Methodology: Y.Z, T.W, M.G, W.L, Z.Z; Project Administration: Y.Z, T.W, W.L, X.W, Z.Z; Resources: T.W, M.G, Z.Z; Supervision: W.L, X.W, Z.Z; Validation: Y.Z, T.W, M.G, W.L; Visualization: Y.Z, T.W, M.W, Z.Z; Writing – Original Draft: Y.Z, T.W, M.W; Writing – Review & Editing: W.L, X.W, Z.Z.

