## Supplementary material for "POI-associated LMD-3 mutation impairs B12-regulated lysosomal function and reproductive capacity in *C. elegans*": 663224_0_supp_11621226_t4488l.pdf

Yile Zhai *et al.*

**This PDF file includes:**

Fig. S1. Sequence alignment highlighting conserved domains and the POI-associated residue.

Fig. S2. The *lmd-3*$^{W690C}$ mutation in *C. elegans* impairs fecundity and increases germline apoptosis

Fig. S3. The W690C mutation impairs protein degradation and lysosomal function.

Fig. S4. meCbl supplementation restores LMD-3(W690C)-impaired autophagy and protein clearance.

Fig. S5. The W690C mutation impairs LMD-3 gene expression and compromises protein stability.

Table S2. Summary of examined reporters and associated germline defects in *lmd-3*$^{W690C}$ mutants.

Table S4. Machine learning-based prediction of LMD-3$^{W690C}$ stability.

**Other Supplementary Materials for this manuscript include the following:**

Movie S1. Structural conservation of the TLDc domains. (Data provided in a separate .mp4 format file)

Table S1. *C. elegans* strains used in this study. (Data provided in a separate Excel spreadsheet)

Table S3. Primers and oligos used in this study. (Data provided in a separate Excel spreadsheet)

Source data 1. Comprehensive record of experimental raw data. (Data provided in a separate Excel spreadsheet)

Source data 2. Uncropped scans of western blot membranes. (Images provided in a separate Word document)

### Supplementary Figures. 1 to 5

### Supplementary Fig. 1

```
LMD-3    0 ---------------------------------------------------------------------- 0
NCOA7    1 MDTKEEKKERKQSYFARLKKKQAKQNAETASAVATRTHTGKEDNNTVVLEPDKCNIAVEEEYMTDEKKK 70
                                 Q23X
                                                                 LysM
LMD-3    0 -------------MGSFGSVSS---------VPPGSAKIPAFQMDYTITETDTLERVAASHDCTVGELMK 48
NCOA7   71 RKSNQLKEIRRTELKRYYSIDDNQNKTHDKKEKKMVVQKPHGTMEYTAGNQDTLNSIALKFNITPNKLVE 140
                        :  : *:..              .: *   *:**  : ***: :* ..: * .:*::
                                                                   A128T

LMD-3   49 LNKMASRMVFPGQKILVPLANADDVFDPPSSASSLKSSGNQISGIENAEDGIRKGPGGAVPAHVRRGSFT 103
NCOA7  141 LNKLFTHTIVPGQVLFVPDANSPSSTLRLSSS----SPGATVSPS-SSDAEYDKLP---------DADLA 196
           ***: :: :.*** ::** **: .     **:    * *  :*   .::    * *          ..::
              T146I
                                             GRAM
LMD-3  104 KTQSAPIPR--GSNDEADTDCLQRFLKIKVKQVTESDGTVSGTLLVTPNCLMFDPDVSHPLVKENGPDLY 186
NCOA7  197 RKALKPIERVLSSTSEEDEPGVVKFLKMNCRYFTDGKGVVGGVMIVTPNNIMFDPHKSDPLVIENGCEEY 266
           :.   ** *  .*..* *   : :***:: : .*:..*.*.*.::**** :****. *.*** *** : *

LMD-3  187 GMVANMDEIQSVSVYKEISGLTGDKAEK------KRDIFDPDHLRTPED----SP--------KKQIKQN 238
NCOA7  267 GLICPMEEVVSIALYNDISHMKIKDALPSDLPQDLCPLYRPGEWEDLASEKDINPFSKFKSINKEKRQQN 336
           *::. *:*: *:::*::** :. ..*           :: *.. .   .    .*        *:: :**

LMD-3  239 GEIEIAKNEEVAFEAGSVDSNHSPKFGSDIALPAISEETKDSPCEEAKGDLSRNRSATLGDEEIQLRAR- 307
NCOA7  337 GEKIMTSDSRPIV-PLEKSTGHTPTKPSGSS------------VSEKLKKLDSSRETSHGSPTVTKLSKE 393
           **  ::.:..  .   . .:.*:*.  *.  :             .*   .*.  .*.:: *.  :   ::

LMD-3  307 -SRTSSQASSN-----DERPRSFSELDTPSEGKNIG-----------RFSPNAARRSFGKLGRTLSARAK 361
NCOA7  394 PSDTSSAFESTAKENFLGEDDDFVDLEELSSQTGGGMHKKDTLKECLSLDPEERKKAESQINNS--AVEM 461
            * ***  .*.       .  .* :*:  *. ..  *            :.*:  ::: .:::..:  *

LMD-3  362 SIQGTVTSGAEKVVGTA----VQGTKTV----AHGVVTHTKSAADTLQSGIEN----------------- 407
NCOA7  462 QVQS-----ALAFLGTENDVELKGALDLETCEKQDIMPEVDKQSGSPESRVENTLNIHEDLDKVKLIEYY 526
           .:*.     *  .:**     ::*:  :     :.:: .... :.: :* :**

LMD-3  407 ------GAKAAADVAGRVVDKGQSLMSESIN------------GVTEIFSVPDFEQ-PAKSPLAMKREQS 457
NCOA7  527 LTKNKEGPQVSENLQKTELSDGKSIEPGGIDITLSSSLSQAGDPITEGNKEPDKTWVKKGEPLPVKLNSS 596
                 * :.: ::    :..*:*:   .*:             :**  . **       .** :* :.*

LMD-3  578 LAKLQDLRRQTAEARENSAKENHAGVFACATSS-D--EAPDLFAAVDEIMERNRIQSNDSTASQSPILPF 523
NCOA7  597 TE-A-NVIKEALDSSLESTLD-----NSCQGAQMDNKSEVQLWLLKRIQVPIEDILP--SKEEKSKTPPM 657
                :: ::: ::  :*: :      :*  :. *  .  :*:      :  : *    *.  .:*   *:

LMD-3  524 YMAVRLTRNKKKKKSSTSSPSYDEDIAFG-NKLKREFWFAVPRNQADNIYHFLLQWSPDKYGLDTTTSTT 587
NCOA7  658 FLCIKVGKPMRKSFATHT---AAMVQQYGKRRKQPEYWFAVPRERVDHLYTFFVQWSPDVYGKDAK---- 720
           ::.::: : :*. ::  :         :* .: : *:******::.*::* *::***** ** *:.

LMD-3  588 EEPSSVMSNDGQDKGFIVLGSNADESLGGEKSTPLFGSSLLNREWELVTVSEMCRRLSLD----QELELP 658
NCOA7  720 ------------EQGFVVVEKEELNMI------DNFFSEPTTKSWEIITVEEAKRRKSTCSYYEDEDEEV 772
                       ::**:*: .:  : :        * *.  .:.**::**.*  ** *      :* *
                       TLDc
LMD-3  659 IPDGAATSQILDELMIRQVMDILPPRAEGYPWVNIYNSEKHGFSLATMYRKMAEFDEDLSPVLLIIRDTK 728
NCOA7  773 LPVLRPHSALLENMHIEQLARRLPARVQGYPWRLAYSTLEHGTSLKTLYRKSASLD---SPVLLVIKDMD 839
           :*     * :*::: *.*:   ** *.:****   *.: :** ** *:*** *.:*   *****:*:*
                                          W804C

LMD-3  729 EHVFGAVVSSAIRPNDHFFGTGDSCLLWRFTGEVPHTRELRQYNWTGDNQYFVNAAKDSLSIGAGSGRNG 798
NCOA7  840 NQIFGAYATHPFKFSDHYYGTGETFLY-TF------SPHFKVFKWSGENSYFINGDISSLELGGGGGRFG 902
           .:::*** .: :: .**::***:: *   *      : .:: ::*:*:*.**:*.  .**.:*.*.** *

LMD-3  799 LWLDADLNHGSSQKCETFDNEPLCGDDQDFIIQFIEAYGFRM 840
NCOA7  903 LWLDADLYHGRSNSCSTFNNDILS-KKEDFIVQDLEVWAFD* 942
           ******* ** *:.*.**:*: *. ..:***:* :*.:.*
```

**Supplementary Fig 1. Sequence alignment highlighting conserved domains and the POI-associated residue.** Amino acid sequence alignments with the Clustal Omega program. The conserved LysM, GRAM, and TLDc domains are indicated by different colored backgrounds. Mutations associated with premature ovarian insufficiency (POI) are highlighted in red. The percentage of identical amino acids is 26.46%. Identical amino acids are marked with an asterisk (*), while strong and weak conservation are indicated with a colon (:) and a period (.), respectively.

**Supplementary Fig. 2**

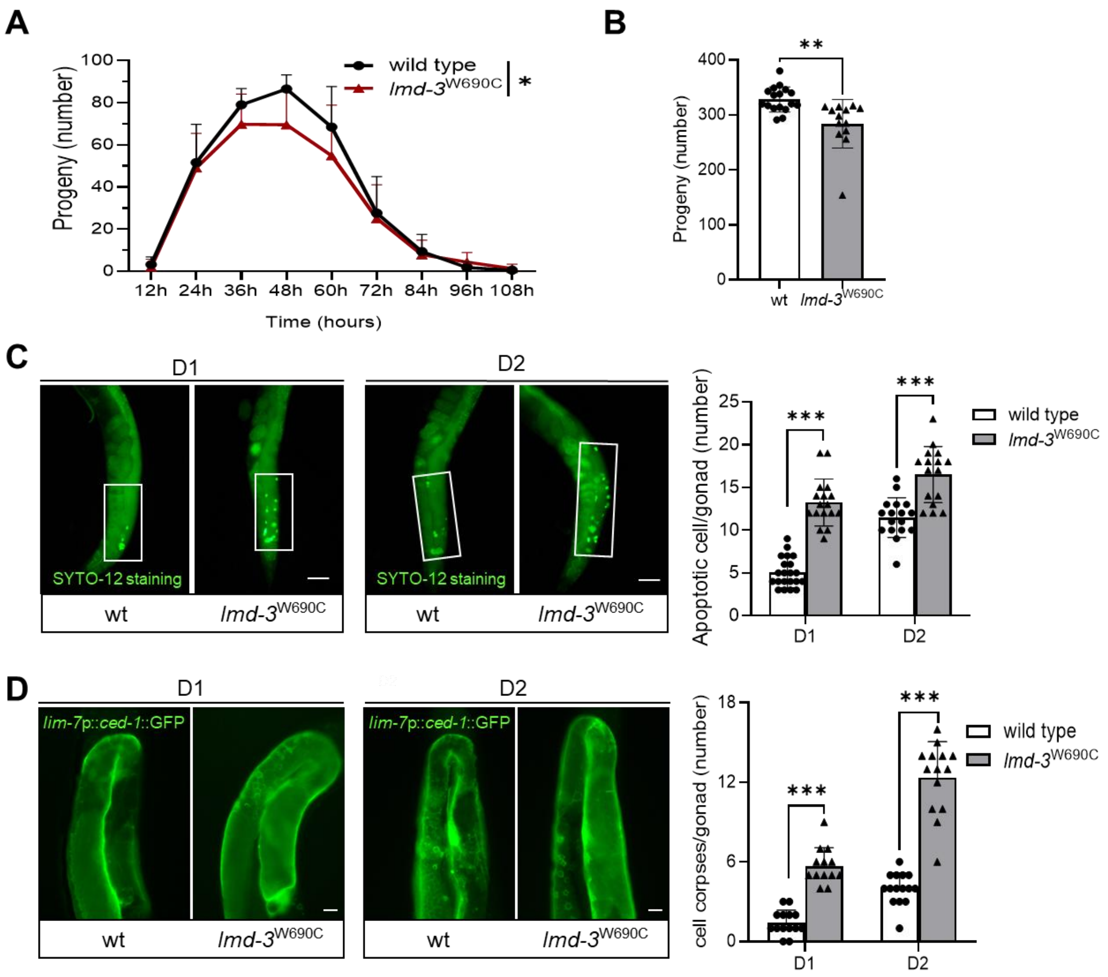


**Supplementary Fig 2. The *lmd-3*$^{\mathrm{W690C}}$ mutation in *C. elegans* impairs fecundity and increases germline apoptosis*.* A** Independent replicates of progeny production curves show the reproductive output of wild-type (black) and *lmd-3*$^{\mathrm{W690C}}$ (red) hermaphrodites (n ≥ 13 per group, *$p < 0.05$). **B** Independent replicates of total brood sizes indicate the overall reproductive ability of wild-type and *lmd-3*$^{\mathrm{W690C}}$ mutants. Data are presented as mean ± SEM (n = 13 per group, **$p < 0.01$). **C, D** Independent replicates of germline apoptosis in wild-type and *lmd-3*$^{\mathrm{W690C}}$ mutants with SYTO12 staining (**C**) and (**D**) a *lim-7p::ced-1::GFP* reporter (n ≥ 13 per group; unpaired t-tests, ***$p < 0.001$). White arrowheads indicate apoptotic cell corpses. The data show a significant increase in apoptosis in the mutants.

D1, Day 1 of adulthood; D2, Day 2 of adulthood. Scale bars, 50 μm.

**Supplementary Fig. 3**

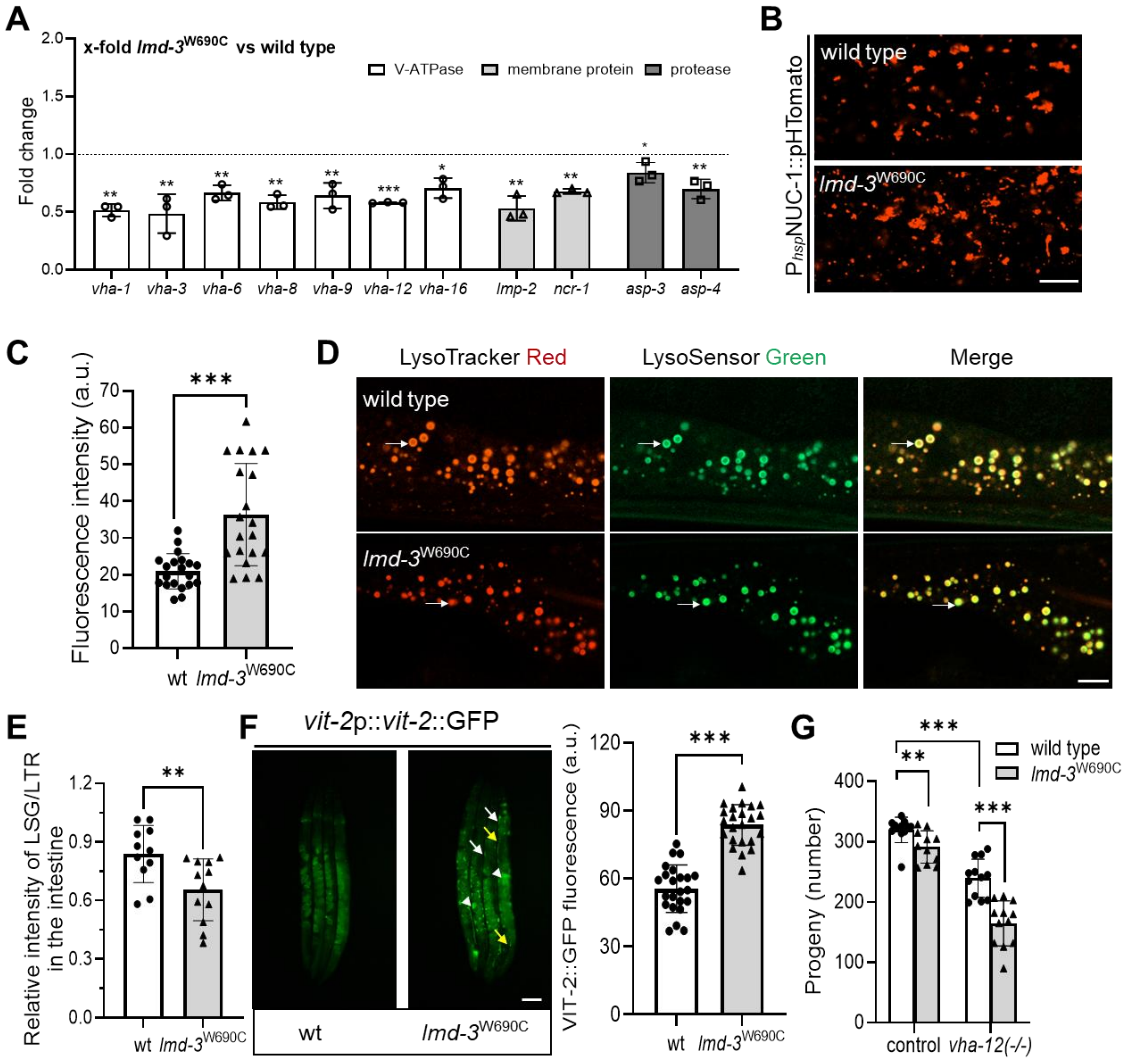


**Supplementary Fig 3. The W690C mutation impairs protein degradation and lysosomal function. A** Independent replicates of transcriptional analysis for lysosome-related genes in wild type and *lmd-3*$^{W690C}$ animals at the D1 adult stage (n = 3 for each group, unpaired t-tests: ***p* < 0.01, ****p* < 0.001, ns, no significant differences). **B, C** Independent replicates of fluorescence images (**B**) and quantification (**C**) of NUC-1::pHTomato expression in the hypodermis driven by a heat-shock (hs) promoter (n ≥ 20 per group, ****p* < 0.001). a.u., arbitrary units. Scale bar, 10 μm. **D, E** Independent replicates of confocal fluorescence images (**D**) and quantification (**E**) of intestines co-stained with Lysosome-Specific Green (LSG DND-189) and Lysosome-Targeting Red (LTR DND-99) in wild type and *lmd-3*$^{W690C}$ animals at the D1 adult stage. White arrowheads indicate vesicular lysosomes stained by both dyes. Relative intensity of LSG/LTR were quantified (n ≥ 11 per group, **$p$ < 0.01). Scale bars, 10 μm. **F** Representative fluorescence images and quantification of VIT-2::GFP accumulation in wild type and *lmd-3*$^{W690C}$ animals at D1 adult stage (n = 23 per group, unpaired t-tests: ns, no significant difference, ***$p$ < 0.001). Aggregates in different tissues: white arrowheads denote germline, white arrows denote pseudocoelom, yellow arrows denote intestines. a.u., arbitrary units. Scale bars, 50 μm. **G** Total brood sizes indicate the effect of *vha-12(ok821)* mutants on reproductive capacity in wild type and *lmd-3*$^{W690C}$ animals (n ≥ 10 per group, **$p$ < 0.01, ***$p$ < 0.001).

### Supplementary Fig. 4

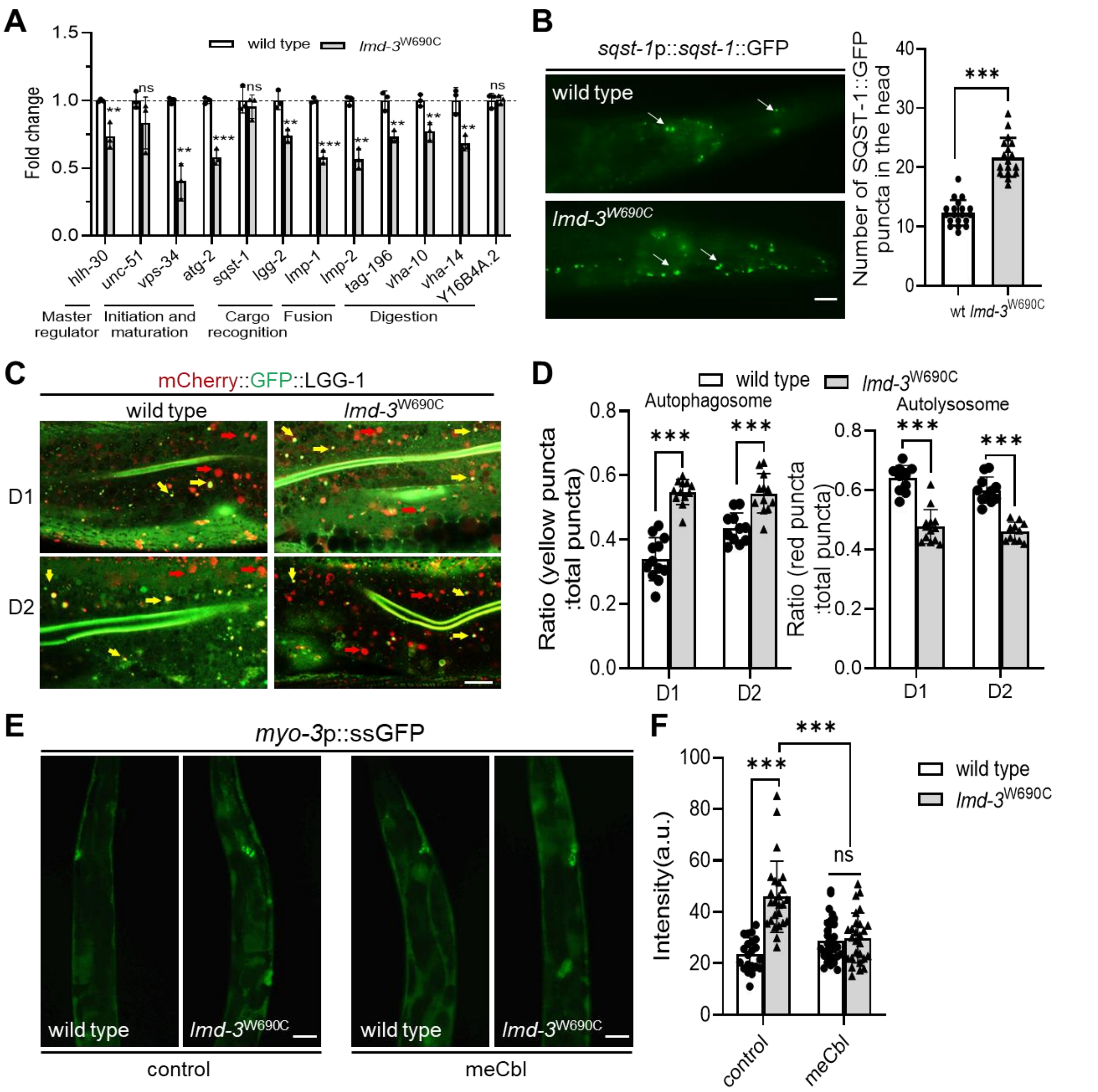


**Supplementary Fig 4. meCbl supplementation restores LMD-3(W690C)-impaired autophagy and protein clearance. A** Independent replicates of transcriptional analysis for key autophagy genes were perfomred wild type and *lmd-3*$^{W690C}$ animals, and gene expression was quantified by RT-qPCR (n = 3 per group, unpaired t-tests: *$p$ < 0.1, **$p$ < 0.01, ***$p$ < 0.001, ns, no significant differences). **B** Independent replicates of fluorescence images and quantification for SQST-1::GFP accumulation in wild type and *lmd-3*$^{W690C}$ animals at D1 adult stage (n ≥ 17 per group, ****p* < 0.001). White arrowheads denote fluorescent protein aggregates. Scale bars, 50 μm. **C, D** Independent replicates of confocal images (**C**) and quantification (**D**) of the mCherry::GFP::LGG-1 reporter in wild type and *lmd-3*$^{W690C}$ animals at D1 and D2 adult stages. The ratio of autophagosomes (yellow puncta, yellow arrow) to total puncta and autolysosomes (red puncta, red arrow) to total puncta was quantified (n ≥ 10 per group, ****p* < 0.001). Scale bar, 10 μm. **E, F** Representative fluorescence images (**E**) and quantification (**F**) of secreted protein accumulation (*myo-3*p::ssGFP) in the coelomocytes of wild type and *lmd-3*$^{W690C}$ animals with mecobalamin (meCbl) supplementation at the D1 stages (mean ± SEM, n ≥ 21 per group, unpaired t-tests: ****p* < 0.001, ns, no significant differences). Scale bars, 50 μm. a.u., arbitrary units.

**Supplementary Fig. 5**

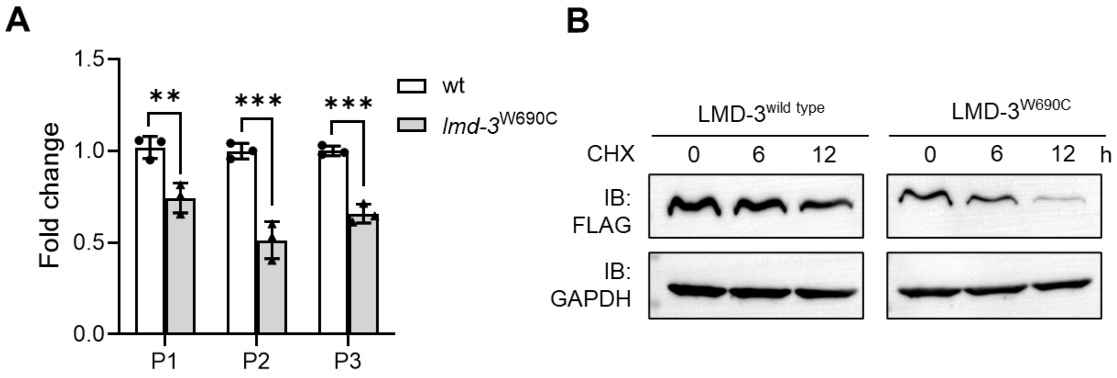


**Supplementary Fig 5. The W690C mutation impairs LMD-3 gene expression and compromises protein stability. A** The independent replicate of qRT-PCR measurements for endogenous *lmd-3* mRNA levels in wild type (wt) and *lmd-3*$^{W690C}$ animals (n = 3 per group, unpaired t-tests: **$p < 0.01$, ***$p < 0.001$)***.*** **B** Independent replicates of Western blot images show the turnover of 1xFLAG-tagged LMD-3 W690C mutation in H293T cells treated with cycloheximide (CHX) for 0, 6, and 12 h.

**Supplementary Table 2**

**Supplementary Table 2. Summary of examined reporters and associated germline defects in *lmd-3*$^{W690C}$ mutants.**

| | *hsp-16.2*p::GFP (UPR$^{cyto}$) intensity | | *hsp-4p::GFP* (UPR$^{ER}$) intensity | | Progeny number | Germline stem cells number | Proliferative zone nuclei number | Meiotic prophase nuclei number |
|---|---|---|---|---|---|---|---|---|
| wild type | L4 | 32.96±3.89 | L4 | 29.98±3.75 | | | | |
| | D1 | 34.26±3.05 | D1 | 33.60±5.77 | 332.59±24.00 | 455.59±13.18 | 182.29±12.37 | 301.43±33.32 |
| | D2 | 34.50±2.63 | D2 | 37.83±6.23 | | | | |
| *lmd-3*$^{W690C}$ | L4 | 31.63±3.31 | L4 | 29.05±5.13 | | | | |
| | D1 | 33.99±2.18 | D1 | 37.25±6.42 | 225.76±73.36 | 418.60±14.26 | 137.14±15.54 | 266.93±30.23 |
| | D2 | 38.93±4.91 | D2 | 44.73±4.62 | | | | |

**Supplementary Table 4**

### Supplementary Table 4. Machine learning-based prediction of LMD-3$^{W690C}$ stability

| Tool Name | ΔΔG (Kcal/mol) | Outcome | Web server |
|---|---|---|---|
| mCSM[a] | -1.458 | Destabilizing | https://biosig.lab.uq.edu.au/duet/ |
| SDM[a] | -1.64 | Destabilizing | https://biosig.lab.uq.edu.au/duet/ |
| DUET[a] | -1.286 | Destabilizing | https://biosig.lab.uq.edu.au/duet/ |
| INPS (sequence)[a] | -1.64 | Destabilizing | http://inpsmd.biocomp.unibo.it. |
| INPS (structure)[a] | -2.15 | Destabilizing | http://inpsmd.biocomp.unibo.it. |
| I-Mutant (sequence)[b] | -1.63 | Destabilizing | https://folding.biofold.org/i-mutant/i-mutant2.0.html |

Six machine learning-based methods were used to predict the change in free energy (ΔΔG) upon W690C mutation. [a], A negative ΔΔG indicates structural destabilization; [b], $\Delta\Delta G < -0.5$ indicates the decrease of stability.

**Movie S1. Structural conservation of the TLDc domains.** (Data provided in a separate .mp4 format file)

**Table S1. *C. elegans* strains used in this study.** (Data provided in a separate Excel spreadsheet)

**Table S3. Primers and oligos used in this study.** (Data provided in a separate Excel spreadsheet)

**Source data 1. Comprehensive record of experimental raw data.** (Data provided in a separate Excel spreadsheet)

**Source data 2. Uncropped scans of western blot membranes.** (Images provided in a separate Word document)
